# Assessment of Phenotype Microarray plates for rapid and high-throughput analysis of collateral sensitivity networks

**DOI:** 10.1101/694109

**Authors:** Elsie J. Dunkley, James D. Chalmers, Stephanie Cho, Thomas J. Finn, Wayne M. Patrick

## Abstract

The crisis of antimicrobial resistance is driving research into the phenomenon of collateral sensitivity. Sometimes, when a bacterium evolves resistance to one antimicrobial, it becomes sensitive to others. In this study, we have investigated the utility of Phenotype Microarray (PM) plates for identifying collateral sensitivities with unprecedented throughput. We assessed the relative resistance/sensitivity phenotypes of nine strains of *Staphylococcus aureus* (two laboratory strains and seven clinical isolates) towards the 72 antimicrobials contained in three PM plates. In general, the PM plates reported on resistance and sensitivity with a high degree of reproducibility. However, a rigorous comparison of PM growth phenotypes with minimum inhibitory concentration (MIC) measurements revealed a trade-off between throughput and accuracy. Small differences in PM growth phenotype did not necessarily correlate with changes in MIC. Thus, we conclude that PM plates are useful for the rapid and high-throughput assessment of large changes in collateral sensitivity phenotypes during the evolution of antimicrobial resistance, but more subtle examples of cross-resistance or collateral sensitivity cannot be reliably identified using this approach.

## INTRODUCTION

Collateral sensitivity is when bacteria develop resistance to one antibiotic, and in doing so increase their susceptibility to one or more others. The phenomenon was first observed in 1952 [1] but it was largely ignored in the subsequent decades. With the emergence of the antimicrobial resistance crisis, collateral sensitivity has garnered new attention because it offers the potential to preserve the utility of our diminishing supply of antibiotics [2-4]. Were collateral sensitivity to be understood in a systematic and predictable manner, a clinician could treat a persistent infection by cycling through antibiotics in such a way that the second antibiotic was chosen for its enhanced efficacy against resistant microorganisms that develop during treatment with the first.

Many studies into the potential of collateral sensitivity have focused on strains of *Escherichia coli* [5-8], *Pseudomonas aeruginosa* [9] or *Staphylococcus aureus* [10, 11] that were subjected to evolution *in vitro*. In each case, a strain was exposed to one antibiotic, resistant mutants were isolated, and their sensitivities towards up to 25 other antibiotics were tested. The clinical relevance of this *in vitro* approach has been questioned because of the stochasticity of evolution [12] and because it emphasises mutationally acquired resistance, rather than the more common scenario of horizontally transferred resistance [13]. In turn, this has led to studies seeking to identify collateral sensitivities that are more directly relevant in clinical settings. For example, 10 genetically-diverse clinical urinary tract isolates of *E. coli* showed broadly conserved patterns of collateral sensitivity to a panel of 16 antimicrobials [14]. With evidence accumulating that collateral sensitivities may indeed be predictable [15, 16] – but with outstanding questions about how to apply this knowledge – the field is primed to advance rapidly.

In this work we aimed to develop, implement and validate a high-throughput screen for collateral sensitivities. Agar dilution and broth dilution methods [17] to determine differences in minimum inhibitory concentration (MIC) are highly accurate but labour intensive. As a result, it is unusual for any given study to test for collateral sensitivities towards >20 antimicrobials. We hypothesised that higher throughput experiments might reveal previously-overlooked sensitivities, which in turn could help to accelerate the field.

Phenotype Microarray (PM) plates have become a widely used tool for phenotypic characterization of microorganisms [18, 19]. Each 96-well PM plate contains different nutrient sources, growth additives or, in the case of PM plates 11-20, antimicrobial compounds. Each of PM plates 11 to 20 contains 24 antimicrobials, present at different concentrations in four wells. A redox dye is added to each well, which changes from colourless to purple in response to microbial metabolic activity. The rate of colour formation can be monitored automatically using an OmniLog instrument; however, we and others have successfully scored colour development by eye [18, 20-22]. Persistent, well-specific variability has also been observed in OmniLog data collected for plates PM 1-10 [23], suggesting that data collection by eye or by OmniLog is equally valid. We set out to assess whether PM plates could provide sensitive and reproducible enough data to be useful in building collateral sensitivity networks. We concentrated on scoring growth data by eye, in order to develop a protocol with the broadest possible applicability.

In our proof-of-principle experiments, we have focused on the Gram-positive bacterium, *S. aureus*. We have compared resistance and sensitivity to 72 antimicrobials (three PM plates) between seven clinical isolates of methicillin-resistant *S. aureus* (MRSA), one laboratory strain, and one descendent of this strain that was evolved *in vitro* towards oxacillin resistance. Overall we found PM plates to provide reproducible data, although their correlation with broth microdilution was more variable.

## MATERIALS AND METHODS

### Materials

Antibiotics and other specialty chemicals were from Sigma-Aldrich (St Louis, MO, USA) unless noted otherwise. Cefazolin, demeclocycline and oxacillin were from Melford Laboratories (Ipswich, Suffolk, UK).

Phenotype Microarray plates (Biolog, Hayward, CA, USA) were used for chemical sensitivity testing. PM plates 11-13 were used as these cover a range of common and clinically relevant antibiotics, as well as other antimicrobial compounds.

### General microbiological methods

*S. aureus* was cultured in Mueller-Hinton (MH) medium (ForMedium, Hunstanton, UK) at 37°C. Seven isolates of community-acquired MRSA were obtained from the collection held at Christchurch Hospital, New Zealand. These were identified as MRSA-BE, -BK, -BR, -SO, -SY, -TT and -UB. The laboratory strain *S. aureus* ATCC 25923 was tested in parallel. Further, *S. aureus* ATCC 25923 was evolved to be more oxacillin resistant by serial passaging. An MH broth culture (1 mL) was incubated at 37°C until the optical density (OD_600_) exceeded 1.0. Then a 10 μL aliquot was used to inoculate a new 1 mL culture with an increased oxacillin concentration. Serial passaging continued until no further bacterial growth was observed and the resulting isolate was termed 25923evo.

### Phenotype microarrays

Cell lines to be assayed were cultured overnight in 3 mL of MH broth. Each saturated culture was diluted to an OD_600_ of 0.5 ± 0.05, for use as the PM plate inoculum as described below.

Each batch of three PM plates was prepared based upon the manufacturer’s protocol for Gram-positive bacteria. Inoculum solution A (6 ml, total volume) was assembled by combining 1 ml of diluted overnight culture with inoculating fluid IF-0 (sold by Biolog at 1.2× concentration) and redox Dye H (sold by Biolog at 100× concentration).

A Gram-positive additive solution (12× concentration) was made, which comprised 24 mM MgCl_2_, 12 mM CaCl_2_, 0.06% (w/v) yeast extract, 0.06% (v/v) Tween 80, 30 mM D-glucose and 60 mM sodium pyruvate.

Inoculum solution B (35 ml, total volume) was prepared by combining the following components: 29.2 ml of inoculating fluid IF-10b (Biolog; sold as a 1.2× stock); 2.92 ml of the Gram-positive additive solution; 350 µl of Dye H (100× stock); and 2.53 ml of inoculum solution A (described above). Aliquots of inoculum solution B (100 µl) were dispensed into each well of PM plates 11-13. Plates were incubated for 24 h at 37°C without shaking.

After incubation, growth in each well was scored by eye. Each well was given a score out of 2: 0 for colourless (no growth); 1 for light purple (intermediate growth); and 2 for dark purple (full growth). Each antimicrobial was present in four wells at increasing concentrations. Growth scores across the four wells were summed. Maximal resistance therefore corresponded to a growth score of 8, whereas a growth score of 0 corresponded to complete sensitivity.

### Determination of minimum inhibitory concentrations

Minimum inhibitory concentrations (MICs) for six antimicrobial compounds were determined by the broth microdilution method, as described previously [17, 20]. The first column of wells in a 96-well plate was filled with 200 µl of MH broth plus the antimicrobial at the maximum concentration to be tested. The remaining wells contained 100 µl MH broth. This was used to begin a two-fold dilution series, which covered 12 concentrations. A 100 µl aliquot from column 1 was transferred into the wells in column 2, pipetted up and down to mix, and repeated for the remaining rows. A 3-µl aliquot of saturated cell culture, diluted to OD_600_ = 0.5, was used to inoculate each well. Plates were sealed with a breathable membrane (Aeraseal) and incubated in an Incumix plate shaker (Select Bioproducts) at 37°C and 600 rpm for 16 h. The percentage of growth inhibition in each well was calculated by measuring OD_600_ in a plate reader (PerkinElmer) and comparing it to a blank and a well with no antibiotic, according to the following formula (Clinical and Laboratory Standards Institute, https://clsi.org/):

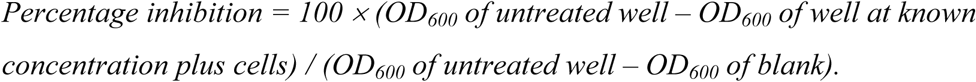

The first well which showed ≥95% inhibition was deemed to be the MIC cut-off for that isolate.

## RESULTS

We assessed the accuracy of PM plates for the discovery of collateral sensitivity networks by assaying seven clinical isolates and two laboratory strains of *S. aureus.* We set out to test two aspects of the accuracy of these assays: reproducibility and reliability. Reproducibility was tested by comparing independent biological replicates in duplicate microarray assays. Reliability was assessed by correlating the relative levels of resistance observed in PM assays with MICs determined using the broth microdilution method [17], which is a particularly common test used in clinical settings [24].

### Reproducibility

All nine strains of *S. aureus* were assayed in duplicate using PM plates 11-13. In total, this experiment therefore probed 648 combinations (9 strains × 72 antimicrobials). The antimicrobials in PM plates 11-13 were grouped according to their mode of action: cell wall-acting antibiotics (16 compounds); protein synthesis inhibitors (22 compounds); inhibitors of nucleic acid synthesis (11 compounds); repurposed cancer and anti-psychotic drugs (7 compounds); and others including antiseptics, disinfectants and metal ions (16 compounds). The full list of compounds is provided in the Supporting Information (S1 Table). As described in the Materials and Methods, relative resistance to each antimicrobial was assessed using a growth score that ranged from 0 (completely sensitive) to 8 (maximally resistant). Independent duplicates were performed several weeks apart. Examples of the PM assays are depicted in Fig 1.

**Fig. 1.**
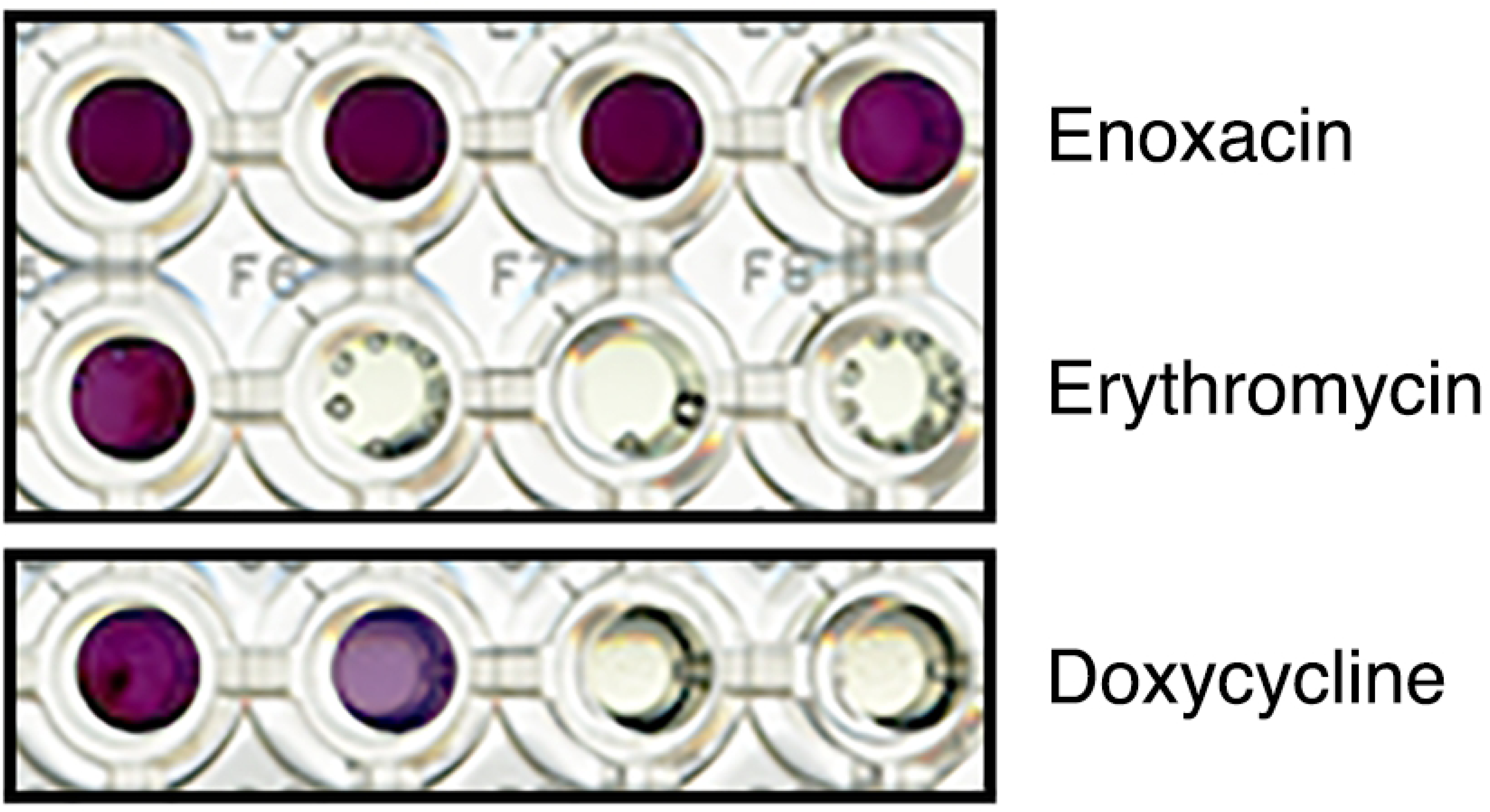
Examples of growth in phenotype microarray plates. The growth of *S. aureus* ATCC 25923 in four wells of increasing enoxacin, erythromycin and doxycycline concentrations is shown. The intensity of colour development (due to the presence of a redox dye) was scored by eye, 24 h after inoculation. It was scored as either 0 (no growth), 1 (intermediate growth) or 2 (full growth). Summing the growth in all four wells yielded a score out of 8 for each antimicrobial. In the examples shown, growths scores of 8, 2 and 3 were recorded for enoxacin, erythromycin and doxycycline, respectively.

Overall, the PM data revealed high levels of resistance in all *S. aureus* strains to many of the compounds tested (Fig 2 and S1-S5 Fig, Supporting Information). As expected, for example, the aminoglycosides were not effective against this Gram-positive bacterium (S2 Fig).

**Fig. 2.**
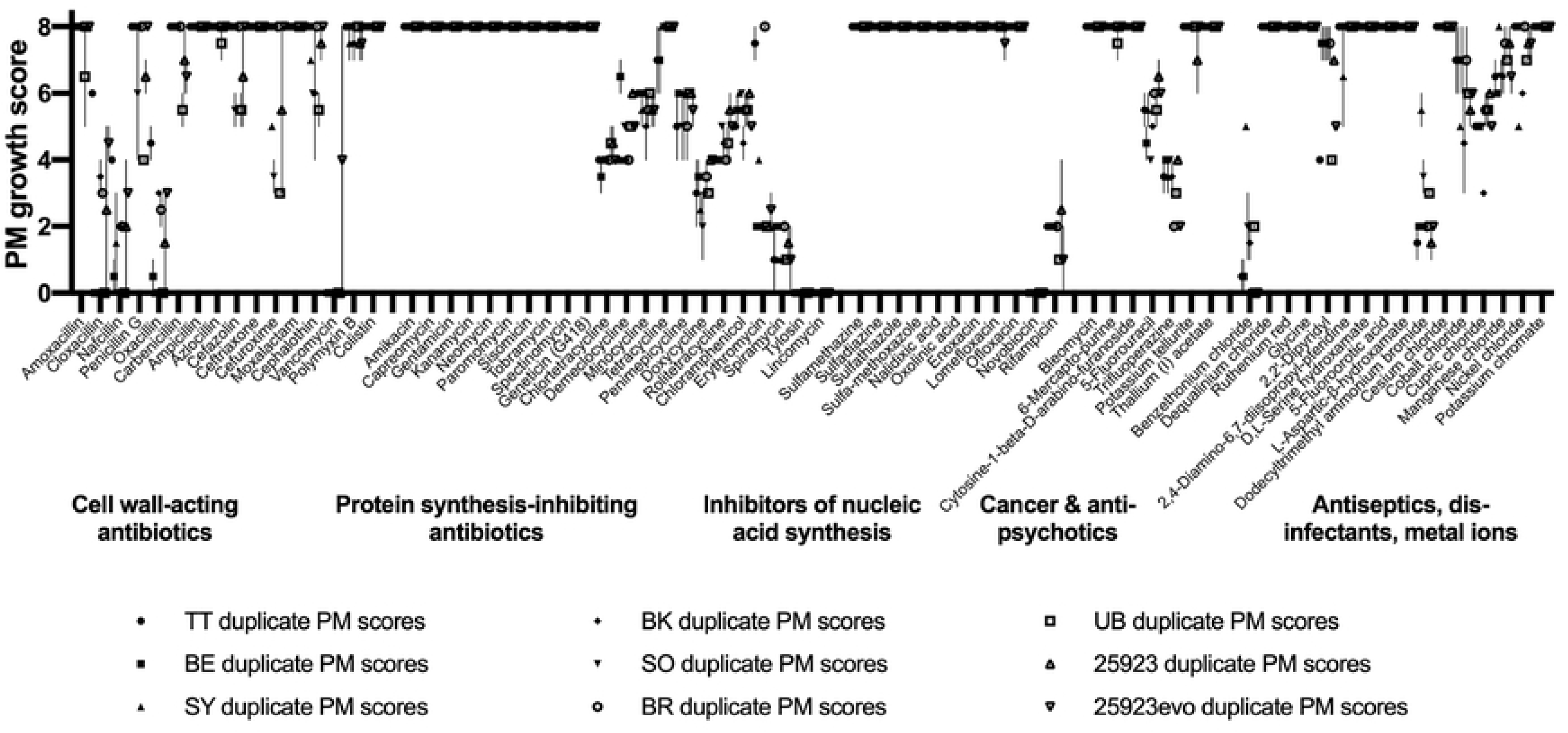
Phenotype microarray plates yield reproducible resistance/sensitivity data. The plot summarizes 1,296 PM growth assays: 72 antimicrobials × 9 *S. aureus* strains, each done in biological duplicate. Each symbol represents the median PM growth score, with the error bars showing the range of the two scores. In 544 of 648 antimicrobial/strain combinations, scores were identical in duplicate assays. In only 29 cases do the error bars span more than one point on the 8-point scale.

Comparing the MRSA isolates with *S. aureus* ATCC 25923 revealed hints of collateral sensitivity; for example, the MRSA isolates all appeared more sensitive to doxycycline (S2 Fig) and 5-fluorouracil (S4 Fig). As expected, *S. aureus* ATCC 25923 showed a low level of oxacillin resistance (average growth score = 1.5), whereas the strain evolved from it in the presence of oxacillin (25923evo) showed increased resistance (growth score = 3).

Comparing the replicated sets of PM data revealed that the growth scores for each antimicrobial were identical between duplicates 84% of the time (Fig 2). When the growth scores were different, it was most commonly only by one point on the 8-point scale. The scores differed by one point 11.5% of the time, and only differed by more than one point in 4.5% of the duplicated assays (29 of 648 combinations). Most of the variability in duplicates was observed for the β-lactams, the tetracyclines and the metal chlorides. In one extreme case, 25923evo returned growth scores of 0 and 8 in duplicate vancomycin tests. The other eight *S. aureus* strains showed reproducible growth scores of 0 for vancomycin, suggesting to us that the score of 8 was the result of a technical or manufacturing error. The overarching conclusion of this experiment was that PM plates yield reproducible data on relative resistance and sensitivity, across both clinical and laboratory strains of *S. aureus.*

### Correlation with broth microdilution

PM plates have been compared favourably against disc diffusion, broth microdilution or molecular methods in terms of throughput, chemical use and time cost [25]. However, the same authors also pointed out that PM plates are best used for preliminary estimates of inhibitory concentrations. We wanted to gauge whether PM assays are sensitive and accurate enough to discover novel collateral sensitivities and build large-scale resistance/sensitivity networks. Therefore, we compared the output from our PM assays with classical MIC determination by broth microdilution.

Within our resistance/sensitivity network (S1-S5 Fig) we noted that the nine *S. aureus* strains responded very similarly to some compounds, but for others the response was highly variable. For example, the clinical isolates MRSA-BR and MRSA-TT were highly resistant to erythromycin (average growth scores of 8 and 7.5 respectively) while most of the other strains we tested were relatively sensitive (growth scores of 2 to 4). On the other hand, all nine strains showed moderate-to-high levels of resistance to cefazolin (growth scores of 5.5 to 8). This reproducible inter-strain variability in the PM data provided us with a way to rigorously assess the degree of correlation with broth microdilution.

We selected six antimicrobial compounds to analyze in more detail. Erythromycin and oxacillin had highly variable efficacies against the nine *S. aureus* strains. Nickel chloride and 2,2’-dipyridyl elicited moderate variability in growth scores between strains. Cefazolin and demeclocycline had relatively uniform efficacies against all strains. The growth scores for each strain against each of these six antimicrobials are summarized in Fig 3. In order to appraise the sensitivity and accuracy of our PM assays, we also determined the MICs for these six compounds. Our goal was to assess whether the large and small growth differences we observed in the wells of PM plates truly reflected differences in antimicrobial resistance/sensitivity.

**Fig 3.**
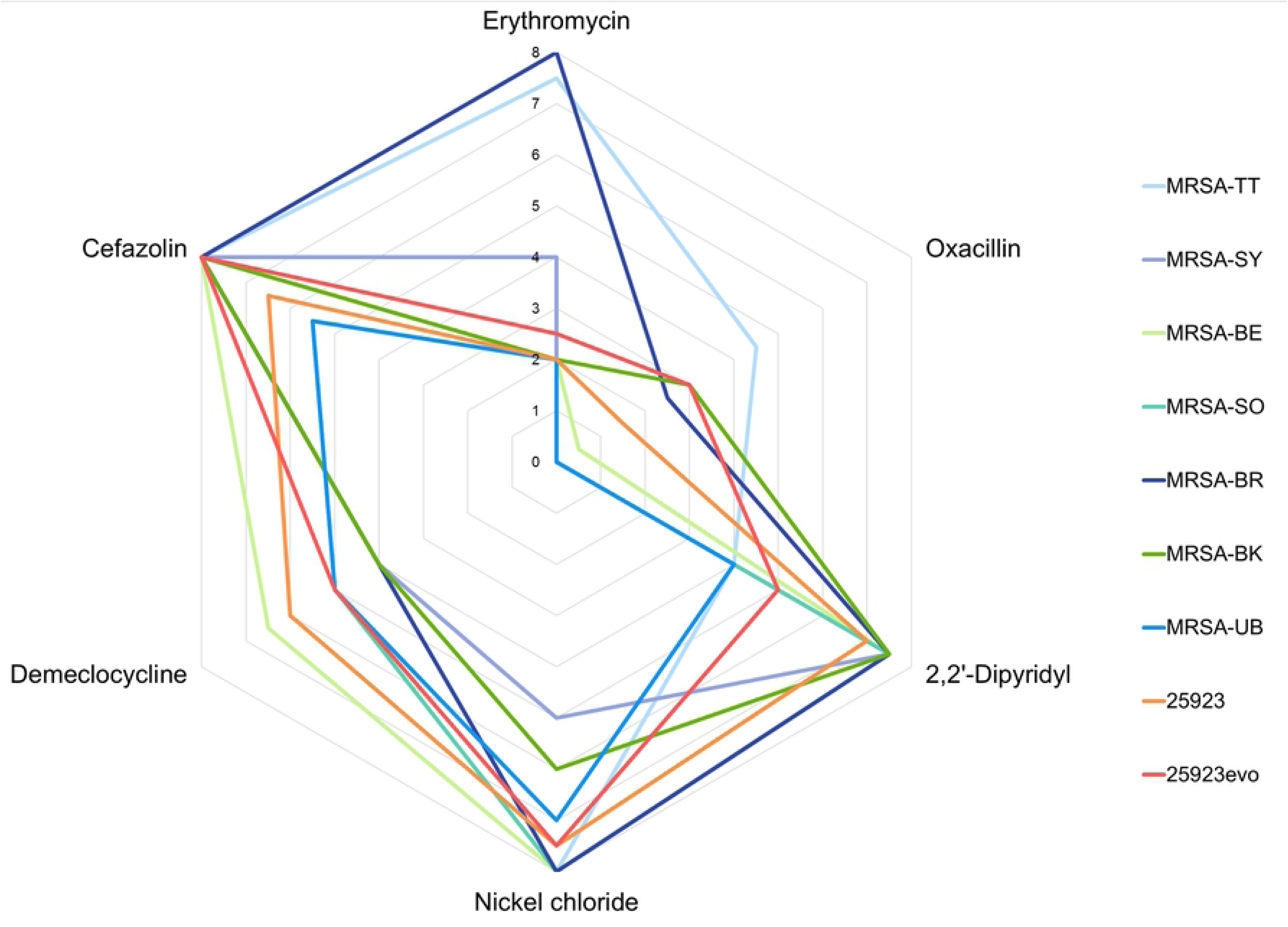
Phenotype microarray growth scores for nine *S. aureus* strains in the presence of six different antimicrobials. The radial axis depicts the average growth score from independent duplicates. The growth scores of the seven MRSA clinical isolates are plotted in shades of blue and green. Data for the laboratory strain *S. aureus* ATCC 25923 and its evolved descendent, 25923evo, are plotted in orange and red.

The comparison of PM growth scores and broth microdilution MICs is shown in Fig 4. In general, large differences in PM growth score faithfully reflected large differences in MIC. For example, a high growth score for erythromycin (8 or 7.5) corresponded to an MIC of 128 µg/ml, whereas a low growth score (2) corresponded to MICs of 8 µg/ml or less. At the same time, the correlation between PM growth score and MIC was far from perfect. The five strains that showed a growth score of 2 in erythromycin-containing PM wells varied in their MICs from 2 µg/ml (MRSA-SO) to 8 µg/ml (MRSA-BE and MRSA-UB). Another strain with an MIC of 8 µg/ml (MRSA-SY) had a growth score of 4, not 2.

**Fig 4.**
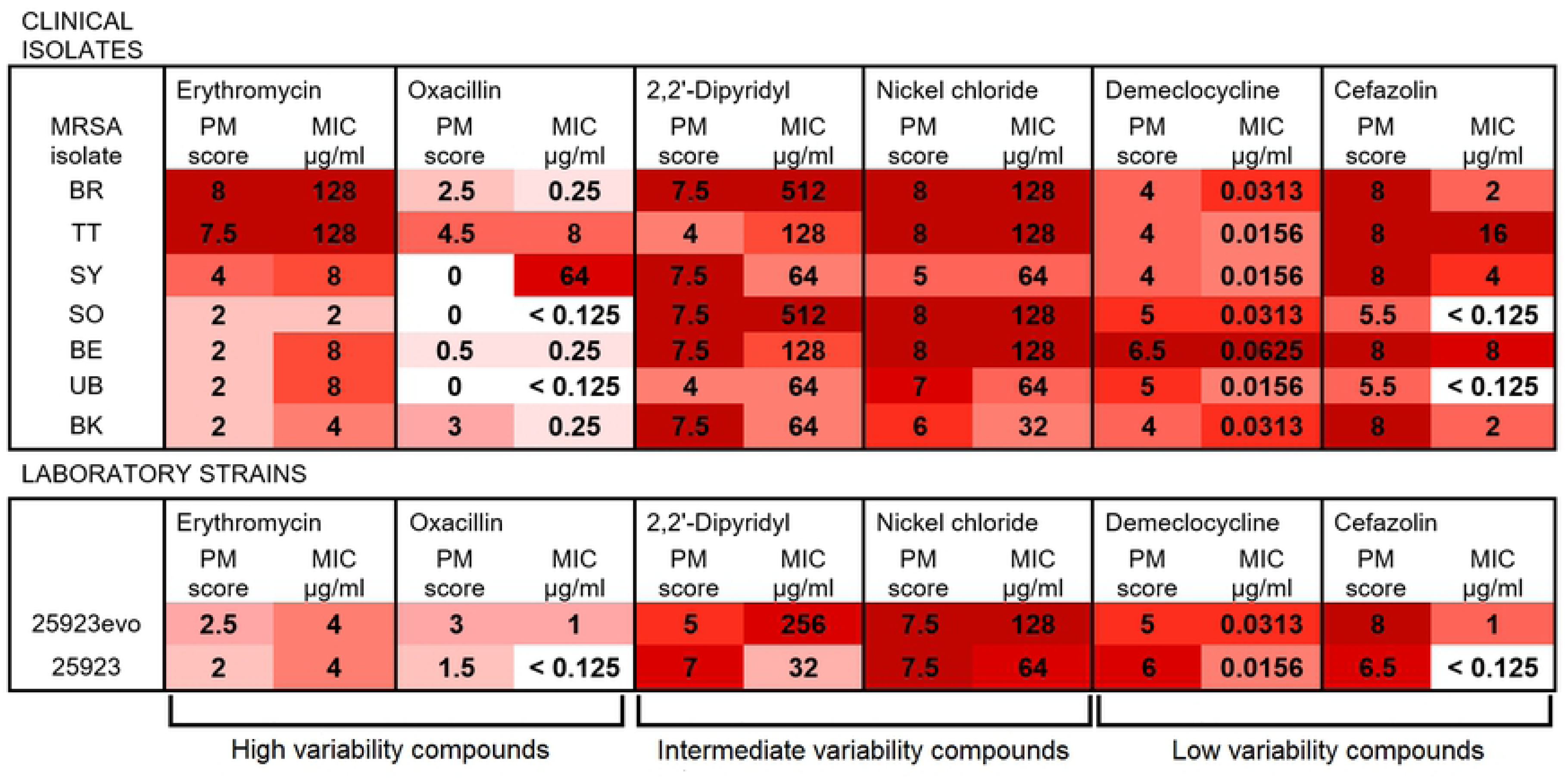
Heat map to correlate PM growth scores and MICs for nine *S. aureus* strains and six antimicrobials. PM growth scores are shaded from dark red (score = 8) to white (score = 0). MICs are also colour-coded, to highlight their level of agreement or disagreement with the corresponding growth score. Compounds are grouped according to the level of variability the nine strains showed in their PM growth scores.

This pattern was consistent across clinical and laboratory strains, and across all of the antimicrobials tested. Results on the polar ends of each scale were more consistent than intermediary scores. For 5 out of the 6 antimicrobials, the highest MICs were directly correlated with the highest PM growth scores (darkest shading in Fig 4). The outlier was MRSA-SY, which reproducibly failed to grow in any oxacillin-containing PM well, but which was highly resistant to oxacillin (MIC = 64 µg/ml) in replicated broth microdilution assays. Similarly, for 5 of the antimicrobials, the lowest MICs were reflected in the lowest PM growth scores (lightest shading in Fig 4). The exception was 2,2’-dipyridyl, for which the most sensitive strain (*S. aureus* ATCC 25923, MIC = 32 µg/ml) grew unusually well in the PM assay (growth score = 7).

On the other hand, differences in PM growth score of less than 2 units could not reliably predict differences in MIC. For example, growth scores of 4, 5 or 6 could all correspond to a demeclocycline MIC of 0.0156 µg/ml (Fig 4). Similarly, growth scores of 5, 7 or even 7.5 could correspond to a nickel chloride MIC of 64 µg/ml.

The manufacturer of PM plates (Biolog Inc.) does not disclose the concentrations of compounds in their plates. The sizes of the incremental increases in concentration across the four PM wells are also unknown for each compound. Not knowing the working range of concentrations complicates the use of PM plates to assess sensitivity/resistance networks. For example, six of the nine *S. aureus* strains showed full growth in all four cefazolin-containing PM wells; that is, they had growth scores of 8. However, the cefazolin MICs of these strains ranged from 1 µg/ml to 16 µg/ml (Fig 4). The most likely explanation is that the four PM wells containing cefazolin range from a very low concentration up to, perhaps, 0.5 µg/ml. If this is the case, any strain with an MIC > 0.5 µg/ml will show full growth in all four wells; however, it becomes impossible to assess the relative resistance or sensitivity of any strain that fulfils this criterion.

## DISCUSSION

This study has emphasized the power – and potential pitfalls – of PM plates for the large-scale assessment of cross resistance and collateral sensitivity. We were able to rapidly obtain resistance and sensitivity data for 72 antimicrobials, which is many more than have been tested in previous studies [8, 12, 14, 15]. It would be straightforward to expand our approach to more of the antimicrobial-containing PM plates. In total, plates 11-20 contain 237 antimicrobials; testing the entire set would represent an order of magnitude increase in screening breadth compared to current approaches. Moreover, scoring PM growth by eye (Fig 1) proved to be a fast, technically straightforward, cost effective and reproducible way to collect resistance and sensitivity data. In independent duplicates, carried out several weeks apart, we obtained identical growth scores in 544 of 648 antimicrobial/strain combinations. Of the remaining combinations, 75 differed by a single point on our 8-point growth scale (Fig 2). This corresponded to the difference between no growth and intermediate growth, or intermediate growth and full growth, in one of the four wells containing a given antimicrobial.

When PM growth scores were carefully compared with MIC data obtained by broth microdilution, a trade-off between throughput and accuracy became apparent. Small differences in PM growth score did not reliably correlate with MIC. Our data suggest that a difference in growth score of at least 2 points is required to indicate a genuine difference in MIC between two *S. aureus* strains. For example, the PM assays suggested that many of the MRSA strains were more sensitive to demeclocyline, nickel chloride or 2,2’-dipyridyl than the laboratory strain *S. aureus* ATCC 25923 (Fig 3). However, the differences in PM growth score were small and the evidence for increased sensitivity was not borne out by MIC testing (Fig 4). The level of agreement (or disagreement) between PM scores and MICs was comparable for the clinical isolates of MRSA and the laboratory strains *S. aureus* 25923 and 25923evo. For the purpose of discovering novel collateral sensitivities, the power of these assays appears to be limited to detecting large reductions in resistance. With one exception (MRSA-SY in oxacillin), differences of ≥16-fold in MIC between any two strains were always correlated with differences in PM growth.

Our results build on previous findings that laboratory strains are not necessarily good models for exploring collateral sensitivity in the clinical setting [12, 26]. Our laboratory-evolved, oxacillin-resistant strain 25923evo frequently behaved closer to its parent, *S. aureus* ATCC 25923, than to the clinical MRSA isolates. Collateral sensitivities could not be extrapolated from 25923evo to the MRSA isolates, either from PM results or from MIC data.

There remains a chasm between the expanding body of laboratory research into collateral sensitivity and the implementation of this research as a therapeutic strategy. A high-throughput assay using PM plates could go some way towards bridging this gap. The assay we have implemented here offers speed and breadth, and is capable of reproducibly identifying large differences in cross-resistance or collateral sensitivity between otherwise closely-related strains.

## ACKNOWLEDGEMENTS

We gratefully acknowledge Prof Stephen Chambers (Director of Infectious Diseases, Christchurch Hospital, New Zealand) and Rosie Greenlees (Canterbury District Health Board) for providing us with the MRSA isolates used in this study. Financial support for this work was received from the Otago Medical Research Foundation, the Maurice and Phyllis Paykel Trust and the Wellington Medical Research Foundation.

## SUPPORTING INFORMATION

**S1 Table. Full list of antimicrobial compounds tested in PM plates 11, 12 and 13, grouped according to mode of action.**

**S1 Fig. Phenotype microarray growth scores for nine *S. aureus* strains in the presence of cell wall-acting antibiotics.** Scores are an average of two biological replicates, with 8 representing maximum relative resistance around the exterior of the radar and 0 representing complete sensitivity at the centre. MRSA isolates are in shades of blue and green. Laboratory strains *S. aureus* ATCC 25923 and 25923evo are orange and red respectively.

**S2 Fig. Phenotype microarray scores for nine *S. aureus* strains in the presence of protein synthesis-inhibiting antibiotics.** Scores are an average of two biological replicates, with 8 representing maximum relative resistance around the exterior of the radar and 0 representing complete sensitivity at the centre. MRSA isolates are in shades of blue and green. Laboratory strains *S. aureus* ATCC 25923 and 25923evo are orange and red respectively.

**S3 Fig. Phenotype microarray scores for nine *S. aureus* strains in the presence of antibiotics that inhibit nucleic acid synthesis.** Scores are an average of two biological replicates, with 8 representing maximum relative resistance around the exterior of the radar and 0 representing complete sensitivity at the centre. MRSA isolates are in shades of blue and green. Laboratory strains *S. aureus* ATCC 25923 and 25923evo are orange and red respectively.

**S4 Fig. Phenotype microarray scores for nine *S. aureus* strains in the presence of repurposed cancer and antipsychotic drugs.** Scores are an average of two biological replicates, with 8 representing maximum relative resistance around the exterior of the radar and 0 representing complete sensitivity at the centre. MRSA isolates are in shades of blue and green. Laboratory strains *S. aureus* ATCC 25923 and 25923evo are orange and red respectively.

**S5 Fig. Phenotype microarray scores for nine *S. aureus* strains in the presence of antiseptics, disinfectants, metal ions, etc.** Scores are an average of two biological replicates, with 8 representing maximum relative resistance around the exterior of the radar and 0 representing complete sensitivity at the centre. MRSA isolates are in shades of blue and green. Laboratory strains *S. aureus* ATCC 25923 and 25923evo are orange and red respectively.

